# IFN-I exacerbates the inflammatory response of epithelial cells to *Chlamydia trachomatis* infection by enhancing TLR3 expression

**DOI:** 10.64898/2025.12.08.692940

**Authors:** Chongfa Tang, Xiaonan Cai, Béatrice Niragire, Félix Louchez, Yaël Victoria Levy-Zauberman, Agathe Subtil, Yongzheng Wu

**Author notes:** Corresponding authors: Agathe Subtil, Institut Pasteur, Université Paris Cité, UMR3691 CNRS, Cellular biology of microbial infection, Paris, France, Yongzheng WU, Institut Pasteur, Université Paris Cité, UMR3691 CNRS, Cellular biology of microbial infection, Paris, France.

## Abstract

The inflammation induced by *Chlamydia trachomatis* infection in the female genital tract (FGT) can have severe consequences. Recent observations in women and mice infected with *Chlamydia* suggest that type I interferon (IFN-I) may have deleterious effects. This study aimed at elucidating the consequences of IFN-I production on *C. trachomatis*-induced inflammation in epithelial cells and the molecular pathway(s) involved. We showed that combination of IFN-I and *Chlamydia* resulted in a stronger induction of inflammation than *Chlamydia* alone, while IFN-I alone had no effect. Inhibiting AKT and mTOR, but not silencing *STAT1*, significantly attenuated the synergetic effect between IFN-I and bacteria on inflammation. Inhibition of ERK also blocked this synergistic effect, although ERK were not activated by IFN-I. We hypothesized that IFN-I enhanced the expression of a pathogen recognition receptor of the host, thereby potentiating detection of the bacteria and the subsequent inflammatory response. IFN-I, but not *C. trachomatis* infection, increased the expression of Toll-like receptor 3 (TLR3), and silencing or knocking-out *TLR3* prevented the synergetic effect between infection and IFN-I. Furthermore, our data support the presence of dsRNA in infected cells and the activation of the MAPK/ERK and AP-1/ATF2 signaling cascades downstream of TLR3. Taken together, our data demonstrate that IFN-I exacerbates the host inflammatory response triggered by *Chlamydia* by increasing TLR3 expression and activation, leading to hyperinflammation. The identified signaling cascades represent potential targets for therapeutical intervention to limit tissue damage upon *Chlamydia* infection.

## INTRODUCTION

Infection by the obligate intracellular bacterium *Chlamydia trachomatis* is the leading cause of sexually transmitted infections (STIs) of bacterial origin (1). The bacteria multiply principally in epithelial cells of the reproductive tract of men and women. In women, the bacteria can ascend from the vagina to the cervix, the endometrium and the fallopian tubes. Epithelial cells from these tissues constitute the first line of defense against *C. trachomatis* infection. In particular, they sense *Chlamydia* through pattern recognition receptors (PRRs) (2–5), distributed either on the cell surface (e.g. TLR2, 4 and 6), in the cytoplasm (e.g. NODs) or in the endosomes (e.g. TLR3, 7, 9) (6). Activation of these PRRs triggers signaling cascades, that eventually lead to expression of genes involved in innate immunity such as pro-inflammatory cytokines and chemokines. Among those, interleukin 6 (IL6) is one of major pro-inflammatory cytokine associated with *Chlamydia* infection (7–9). Inflammation contributes to the resolution of infection. However, in some women, it can result in tissue damage and scar formation, eventually causing severe consequences, such as pelvic inflammatory disease and tubal infertility.

Type I Interferon (IFN-I) belongs to a group of cytokines, that activates several signaling cascades, including Janus kinase 1 (JAK1)/tyrosine kinase 2 (TYK2) and downstream signal transducer and activator of transcription 1 (STAT1), mitogen-activated protein kinase (MAPK) kinase/Erk, phosphoinositide 3-kinase (PI3K)/AKT/p38 and PI3K/AKT/mammalian target of rapamycin (mTOR). These signaling cascades converge towards the induction of expression of interferon-stimulated genes(10). The best-known members of this family, IFNα and β, exert important roles in infectious diseases. While their protective properties against viral infections are well documented, the consequences of their production during bacterial infections are less studied (11, 12). IFN-I production appears to be beneficial to the host during infections by *Streptococcus pyogenes* (13), *Legionella pneumophila* (14, 15), *Bacillus anthracis* (16), *Shigella flexneri* and *Salmonella typhimurium* (17, 18).

IFN-I is synthesized by many cell types, including immune cells. It is secreted by epithelial cells of the female genital tract (FGT) upon *C. trachomatis* infection both in humans and mice, including in an organoid model of infection (19–23). Early works have reported an inhibitory effect of IFN-I on *C. trachomatis* and *C. pneumoniae* replication in epithelial cells (24, 25). However, mice with IFN receptor gene knocked out (IFNAR-KO mice) or deficient for IFN-I signaling cleared *C. muridarum* infection earlier than control mice and developed less oviduct pathology(26). In addition, anti-IFN-I neutralizing antibodies also inhibited bacterial replication and subsequent tissue damage in the FGT of *C. muridarum*-infected mice (22). These data indicate that IFN-I production favors bacterial proliferation and tissue damage in a mouse model of *Chlamydia* infection. Furthermore, retrospective studies showed that both ascending *C. trachomatis* infection in the FGT of patients and *Chlamydia* infection in patients with pelvic inflammatory disease, correlated with increased levels of interferon-inducible protein 10 (IP10), a cytokine produced in response to IFN-I exposure (27, 28). We also observed increased IP10 production by primary epithelial cells exposed to *C. trachomatis* infection (9).

The potential influence of IFN-I on the outcome of *C. trachomatis* infection prompted us to characterize its effect on epithelial cells, which constitute the first sentinels alerting infection of the genital tract. The production of IL6 was markedly enhanced by exposure to IFN-I. By characterizing the molecular mechanism underlying the synergetic effect between IFN-I and *C. trachomatis* infection, we uncovered the central part played by the PRR TLR3.

## MATERIALS AND METHODS

### Cells

The human cervical epithelial cell line HeLa was from ATCC (Virginia, USA). The cells were grown in Dulbecco’s Modified Eagle’s Medium (DMEM) supplemented with glutaMAX^®^ (Gibco, France) and 10% heat-inactivated fetal calf serum (FCS).

Primary epithelial cells were isolated from ecto-cervical explants of patients undergoing hysterectomy as described previously (9). The cells were cultured in keratinocyte serum-free media supplemented with 5 µg/L of human recombinant epidermal growth factor (EGF) and 50 mg/L bovine pituitary extract (BPE) (K-SFM, Thermo Fisher Scientific, #10144892).

### Preparation of TLR3-KO cell line

TLR3-KO HeLa cells were established as described (29). In brief, 0.5 μg of pSpCas9(BB)-2A-Puro (PX459) V2.0 plasmid (Addgene #62988) containing guide RNA (gRNA) duplex sequence of human TLR3 (F: 5’-CACCGTACCAGCCG CCAACTTCACA-3’; R: 5’-AAACTGTGAAGTTGGCGGCTGGTAC-3’) inserted with BbsI enzyme site, was transfected into pre-adhered HeLa cells (0.15 × 10^6^ cells/well, 24-well plate) using 1 μl of jetPRIM^®^ (Polyplus). Four hours later, fresh medium was replaced containing puromycin (2 μg/ml) and cells were incubated for 24h. The surviving cells were detached, counted and serially diluted to collect individual clones in 96-well plates.

To detect the Indel mutation of TLR3, genomic DNA was extracted after expanding the individual clones, using DNeasy Blood & Tissue Kit (Qiagen). The gRNA-containing region (∼ 700 bp) of TLR3 gene was amplified (95°C for 30s, 55°C for 30s, 72°C for 30s, × 35 cycles) from genomic DNA using GoTaq^®^ qPCR system (Promega) and T100^™^ Thermal Cycler (Bio-Rad). The primers for PCR were shown in Table S1 and the primer for sequencing was 5’-GACTTTTGTCACGACTTCAC-3’. Analysis of the sequence using Tracking of Indels by Decomposition (TIDE, https://tide.nki.nl/) revealed the presence of 5 bp (2 alleles) and 6 bp (1 allele) deletions, consistent with the presence 3 copies of the TLR3-containing chromosome 4 in HeLa cells (30).

### Bacterial preparation and infection

*C. trachomatis* LGV-L2 (434/Bu/ATCC) and LGV-L2^IncD^mCherry (stably expressing the fluorescent protein mCherry) (31) strains were propagated in HeLa cells and purified as described (32). Purified elementary bodies (EBs) were stored in 220 mM sucrose, 10 mM sodium phosphate [8 mM Na_2_HPO_4_-2 mM NaH_2_PO_4_], 0.50 mM L-glutamic acid) (SPG buffer) at-80°C. For infection, adhered cells were incubated with bacteria suspended in DMEM containing 10% FCS for different times (depending on the experiments) before harvesting the samples. Unless otherwise specified the infections in Hela cells were conducted with multiplicity of infection (MOI) of 2, and five times more bacteria were used in experiments with primary cells (9), to reach >85% of cells being infected.

### Flow cytometry

Cells (0.3×10^6^ cells/ml) were seeded in 12-well plates. Twenty-four hours post IFN-I (Biogen Idec) treatment (2.5 ng/ml) and/or *Chlamydia* infection, the cells were incubated with 5 μg/ml brefeldin A solution (Biolegend #420601) for 6h to block cytokine secretion. In certain experiments, the cells were pre-treated with pharmacological inhibitors for 1 h before bacterial infection and/or IFN-I treatment. Cells were then detached with 0.5 mM EDTA in PBS, fixed by 2% PFA/2% sucrose (w/v) in PBS for 20 min at room temperature, followed with quenching by 50 mM NH_4_Cl in PBS for 10 min. The cells were then permeabilized with 0.3% (v/v) Triton X-100 in PBS for 10 min and blocked in 1% BSA in PBS for 1h, following with incubation in PBS, 0.1 % BSA with or without 1/40 dilution of PE/Cyanine7-conjugated anti-human IL6 (Biolegend, #501119) for 1h. After washes, the cells were resuspended in PBS and analyzed by CytoFLEX flow cytometer (Beckman Coulter). The data were analyzed using FlowJo (version 10.0.7).

### siRNA treatment

One and half microliter siRNA (10 μM stock) was mixed with 0.75 μl Lipofectamine iMax (Invitrogen) in 50 μl of DMEM media (Gibco®) for 10 min at room temperature. The mixture was added into culture dish before adding cells (0.15×10^6^ cells/well for 24-well plate) suspended in 0.45 ml DMEM/FCS medium, mixed and incubated for 24 h before treatment or infection.

### Immunofluorescence

HeLa cells, p65-GFP expressing HeLa cells (33) were seeded on coverslips in a 24-well culture dish (0.15×10^6^ cells/well). The cells were treated with recombinant human IL1β (10 ng/ml, Thermo Fisher Scientific), IFNβ (2.5 ng/ml), and/or infected with *Chlamydia* strains (MOI=1) as described in the legends of the figures. The cells were then fixed with 4% PFA and 4% sucrose for 20 min at room temperature, followed with quenching with 50 mM NH_4_Cl in PBS for 10 min. The cells were then permeabilized with 0.3% (v/v) Triton X-100, 0.1% (w/v) BSA. Inclusions were stained using rabbit antibodies against the inclusion protein Cap1 (obtained by the lab) and secondary anti-rabbit Alexa Fluor647 antibody, DNA was stained using 0.5 µg/ml Hoechst 33342

(Thermo Fisher Scientific) in PBS containing 0.1 % of BSA. dsRNA was detected using primary monoclonal J2 antibody (Scicons, #RNT-SCI-10010200) at dilution of 1/50 and secondary anti-mouse Alexa Fluor488 antibody. A duplicate infected coverslip was treated with 100 U/ml RNAse-III (Ambion, AM2290) in PBS for 30 min with gentle agitation (20 rpm/min) at room temperature after fixation(34), followed by immunostaining. Coverslips were mounted on slides in a Mowiol solution. Images were acquired on an Axio observer Z1 microscope equipped with an ApoTome module (Zeiss, Germany) and a 63× Apochromat lens. Images were taken with an ORCA-flash4.0 LT camera (Hamamatsu, Japan) using the software Zen.

### Quantification of dsRNA fluorescent signals

All segmentations were performed using Cellpose (35) pretrained model “cyto” with “diameter” parameter set to “0”. Cells were segmented based on the green/dsRNA staining and uv/DAPI channels. Inclusions were segmented based on the red/mCherry channel. Cell nuclei and inclusions appeared in the uv/DAPI channel (DNA stain). Segmentation of this channel using Cellpose resulted in masks corresponding both to inclusions and cells nuclei. The different steps described below were automatically performed using a Fiji macro script (36). Regions of interest (ROI) corresponding to inclusions and cells were respectively defined using the inclusion and cell masks from Cellpose. For each cell ROI, the macro checked each inclusion ROI one by one to see if there was an overlap with the cell ROI. This was done by selecting the intersection of both ROI (“AND” function of the ROI Manager) and by checking if it was not empty (selection Type >-1). Cells were considered infected if they overlapped with at least one inclusion. Then areas of each cell corresponding to nuclei or inclusions were set to intensity values of 0 by subtracting the DAPI and inclusions masks from Cellpose.

The total intensity in the cytoplasm in the dsRNA channel was then measured for each cell. Finally, for each field, two composite pictures of DAPI, mCherry and dsRNA channels were saved, displaying either infected or non-infected cells segmentation for postprocessing verification (see Fig. S6). Finally, RStudio software was used to plot the dsRNA total fluorescence intensity and performed Student’s t-test using ggplot2, ggprism, ggbeeswarm and ggpubr packages.

### RT-PCR, PCR and quantitative PCR

Adhered cells (0.15×10^6^ cells/ml) in 24-well plate were treated with the indicated concentration of recombinant IFN-I, or infected with *C. trachomatis*. Total RNAs were isolated 24 h post IFN-I treatment or 40 hours post infection (hpi) with the RNeasy Mini Kit (Qiagen) and RNA concentrations were determined with a spectrophotometer NanoDrop (Thermo Fisher Scientific). Reverse transcription (RT) was performed using the M-MLV Reverse Transcriptase (Promega) and quantitative PCR (qPCR) was undertaken on the complementary DNA (cDNA) with LightCycler 480 system using SYBR Green Master I (Roche). Data were analyzed using the 2^-^β^ΔΔCt^ method with the *actin* gene as a control gene and results were presented as relative quantity (RQ) compared to untreated or uninfected control cells.

The primers used are listed in Table S1.

### Western blot

Cells were lysed in urea buffer (30 mM Tris, 150 mM NaCl, 8 M urea, 1 % SDS, pH=8.0) containing protease and phosphatase inhibitors. Equal volumes of cell lysates were subjected to SDS-PAGE, transferred to polyvinylidene difluoride (PVDF) membranes and immunoblotted with the proper primary antibodies diluted in PBS containing 5% (w/v) fat-free milk and 0.01% (v/v) Tween-20. Primary antibodies used were rabbit anti-human STAT1 (#9172), Erk (#4695), p38 (#9212), AKT (#9272), rabbit anti-human phosphorylated STAT1 (#9167), Erk (#4370), p38 (#9215), AKT (#9275) and mTOR (#2971), which were purchased from Cell Signaling, rabbit antibodies against the heat shock protein 60 of *Chlamydia* (obtained by the lab), and mouse anti-human β-actin (Sigma, #A5441). Immunoblots were analyzed using horseradish peroxidase secondary antibodies (Abliance), and chemiluminescence was analyzed on a ChemiDoc Touch Imaging System (Bio-Rad).

### Statistical analysis

The experimental data were analyzed using Prism10 (GraphPad). Unpaired t-tests was used for two group comparisons.

## RESULTS

### IFN-I enhances the inflammatory response to C. trachomatis infection

Primary ecto-cervical epithelial cells were infected with *C. trachomatis*, in the presence or absence of recombinant human IFNβ, and transcripts level for *IL6* were measured 30 hpi. The transcription of *IL6* triggered by *Chlamydia* infection was enhanced by IFNβ treatment, when IFNβ alone had no effect (Fig. 1A). To test whether this increase in transcription correlated with higher cytokine levels we used flow cytometry to detect IL6. This experiment required more cells and was therefore conducted in the cervical cell line HeLa. Cells were infected for 30 h in the presence or not of IFNβ. In the last 6 h of infection 5 mg/ml brefeldin A (BFA) was added to the culture medium to block protein secretion. After fixation and permeabilization IL6 was stained with antibodies coupled to a fluorochrome and samples were analyzed by flow cytometry. Addition of IFNβ increased the percentage of cells positive for IL6, compared to infection alone (Fig. 1B, 1C). Consistent with the data obtained in primary cells, IFNβ alone did not elicit IL6 production in HeLa cells (Fig. 1B, 1C). Similar results were obtained using IFNα (Fig. S1A), and the response was dose-dependent (Fig. S1B). These data suggested a synergetic effect of IFN-I on the *Chlamydia*-induced inflammation in primary epithelial cells as well as in HeLa cells. IFNβ was used in the rest of the study.

**Fig. 1.**
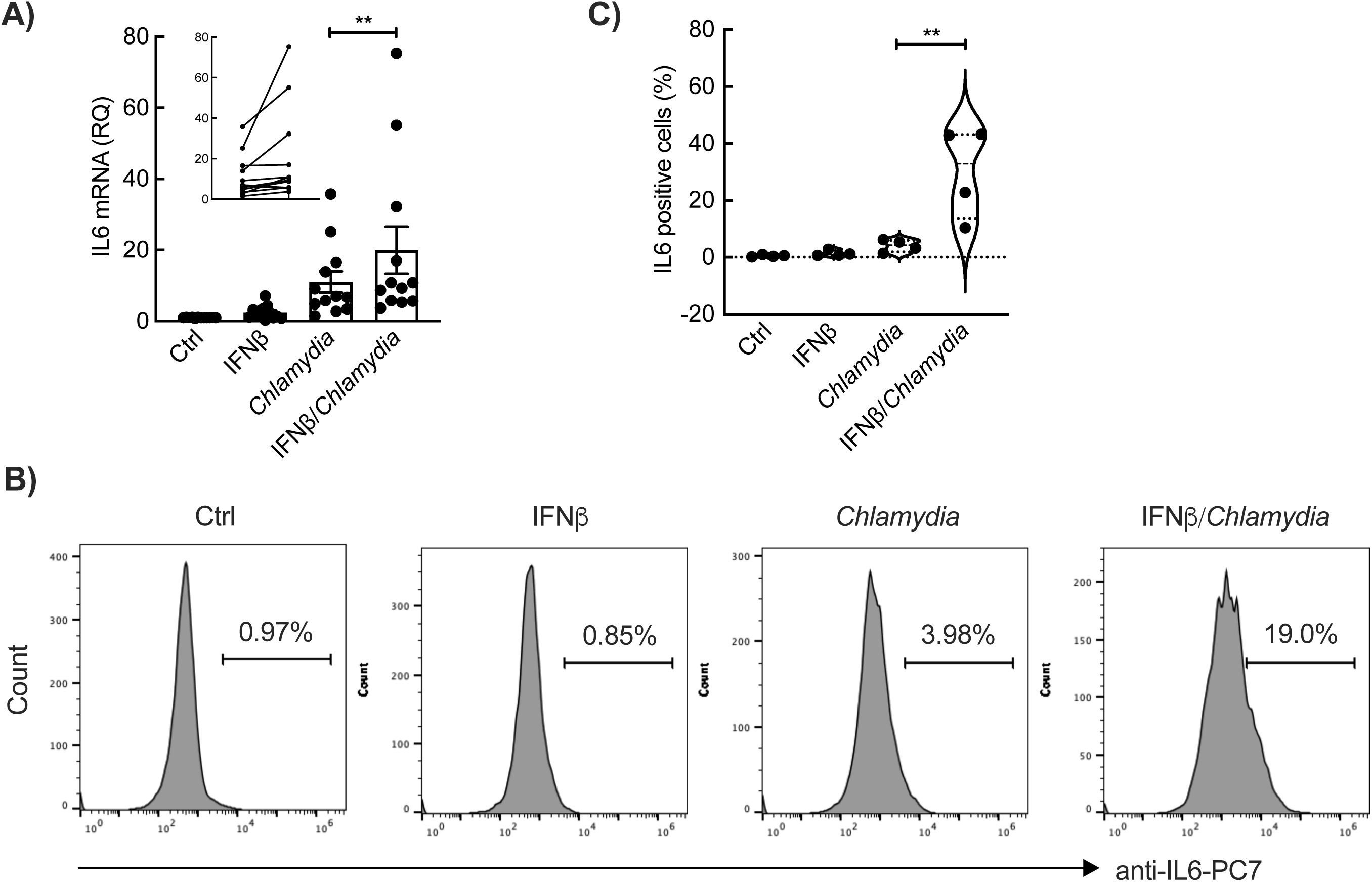
IFNβ enhances the inflammatory response of epithelial cells to *C. trachomatis* infection. (A) Primary cervical epithelial cells were incubated with IFNβ for 24 h, or with *C. trachomatis* for 40 h in the absence or presence of IFNβ. The expression of inflammatory cytokines IL6 were detected by quantitative PCR. IL6 transcript was measured by real-time RT-qPCR and normalized to actin transcript following the 2^-ΔΔCt^ method. The data are presented as relative mRNA levels compared to uninfected cells, five independent experiments are show, each measurement is done in triplicate, and the dots represent different individuals. The insert with paired data displays the *IL6* transcripts in infected cells with or without IFNβ treatment. (B-C) HeLa cells were incubated with IFNβ and/or *C. trachomatis* for 24 h prior to adding brefeldin A for 6 h. After the treatment, the intracellular level of IL6 was measured by flow cytometry using anti-human IL6-PC7 antibody. The histograms are from one representative experiment (B) and the quantification from 4 experiments is presented as the violin plot (C). P-values of Student’s paired t-test are shown when significant (** for p-value < 0.01).

### The synergy between IFN-I and Chlamydia infection requires TLR3 expression

Host cells sense invading microorganisms by recognition of pathogen associated molecular patterns (PAMPs), via specific PRRs. We hypothesized that IFN-I might facilitate the detection of the bacteria and thereby enhance the inflammatory response to the infection. To test this hypothesis, we measured the transcription of the genes coding for the PRRs TLR2/3/4, that were previously implicated in the detection of *C. trachomatis* (2–5). IFN-I stimulated the expression of TLR2, TLR3 and TLR4 by primary epithelial cells both in non-infected and infected cells (Fig. 2A). Similar results were observed in HeLa cells (Fig. S2A). Thus, the three PPR tested could potentially be implicated in the synergetic effect between IFN-I and infection. However, silencing *TLR2* and *TLR4* expression did not affect the level of IL6 production upon *Chlamydia* infection in the presence of IFN-I in HeLa cells (Fig. S2B), while silencing *TLR3* (Fig. 2B, S2B) did, indicating that TLR3 was required. We next generated a *TLR3* KO clone in HeLa cell (Fig. S2C). The production of IL6 in *TLR3* KO cells upon *Chlamydia* infection in the presence of IFNβ was low compared to wild type cells (Fig. 2C), confirming that the synergy between IFN-I and infection implicated TLR3.

**Fig. 2.**
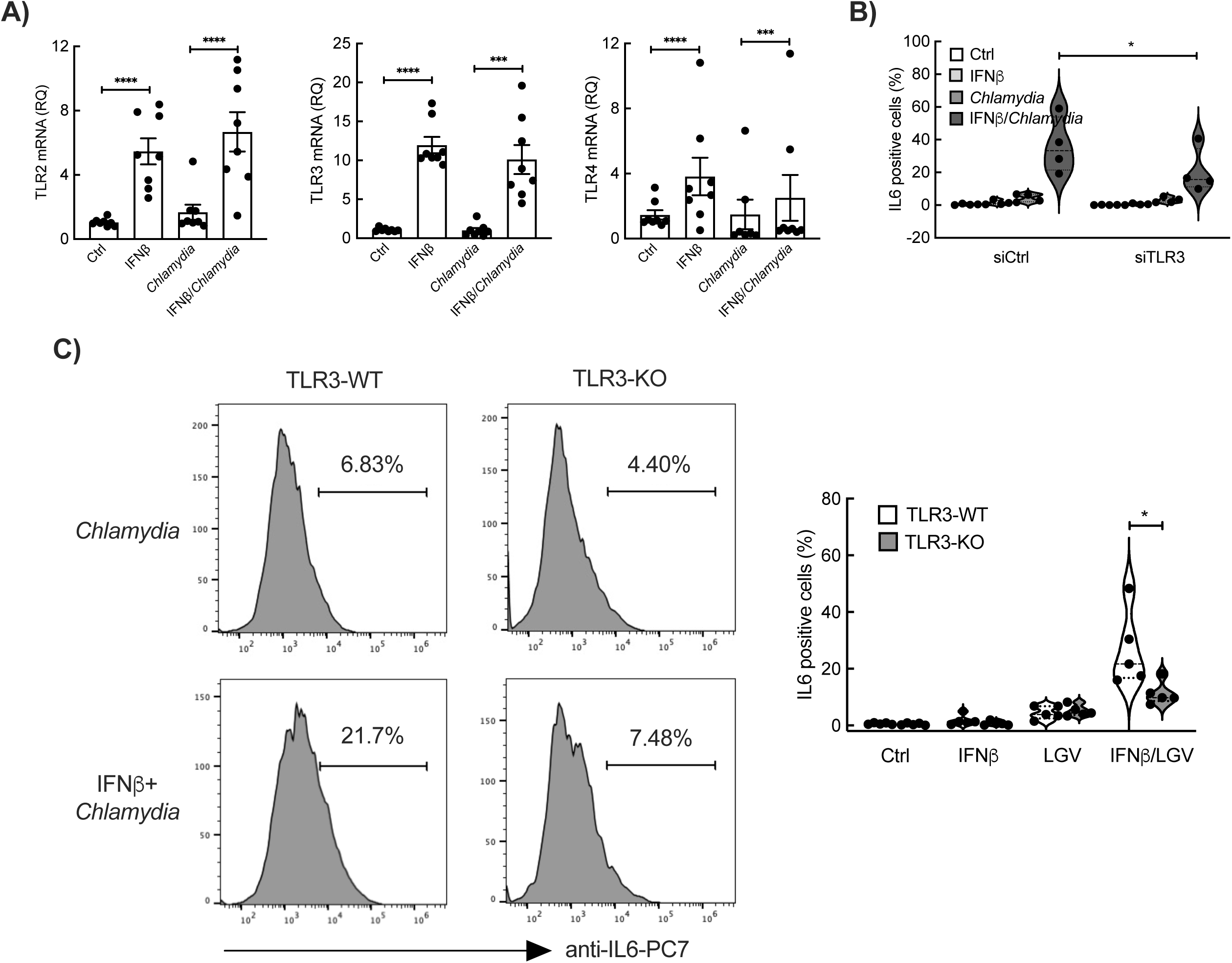
TLR3 mediates the synergy process between IFNβ and *C. trachomatis.* (A) Primary cervical epithelial cells were incubated with IFNβ and/or *C. trachomatis*. Twenty-four hours later, the expression of pattern recognition receptors (PRRs) was measured by real-time RT-qPCR and normalized to actin transcript following the 2^-ΔΔCt^ method. The data are presented as relative mRNA levels compared to untreated cells and shown as the mean±SE with individual values of three independent experiments. Each experiment was conducted in technical triplicates and each dot represented the data obtained with cells isolated from one individual. P-values of Student’s unpaired t-test are shown (*** for p < 0.001, **** for p <0.0001). (B) siRNA was incubated with HeLa cells for 24 h before treatment with IFNβ and/or *C. trachomatis*. Brefeldin A was added 24 hpi for 6 h. Intracellular IL6 protein was measured by flow cytometry using anti-human IL6-PC7 antibody. Data from 4 independent experiments are shown. P-values of Student’s unpaired t-test are shown (* for p < 0.05). (C) TLR3-KO cells or parental HeLa cells were infected with *C. trachomatis* in the presence or absence of IFNβ for 30 h before measuring intracellular IL6 levels by flow cytometry. The histograms are from one representative experiment and the quantification of five independent experiments is presented as mean±SE (right panel). P-values of Student’s unpaired t-test are shown (* for p < 0.05).

### The synergy between IFN-I and infection requires the PI3K/AKT/mTOR but not the STAT1 signaling pathways

Which of the IFN-I activated signaling cascades contributed to the synergy between IFN-I and *C. trachomatis* infection? We first looked at the expression level and phosphorylation of the protein signal transducer and activator of transcription 1 (STAT1). Expression and phosphorylation of STAT1 were increased in cells treated with IFNβ, but not when the cells were infected (Fig. 3A). Moreover, silencing STAT1 had no effect on IL6 production by infected cells treated with IFN-I (Fig. 3B). These data suggest that STAT1 does not contribute to the synergy between IFN-I and *C. trachomatis* on the pro-inflammatory response.

**Fig. 3.**
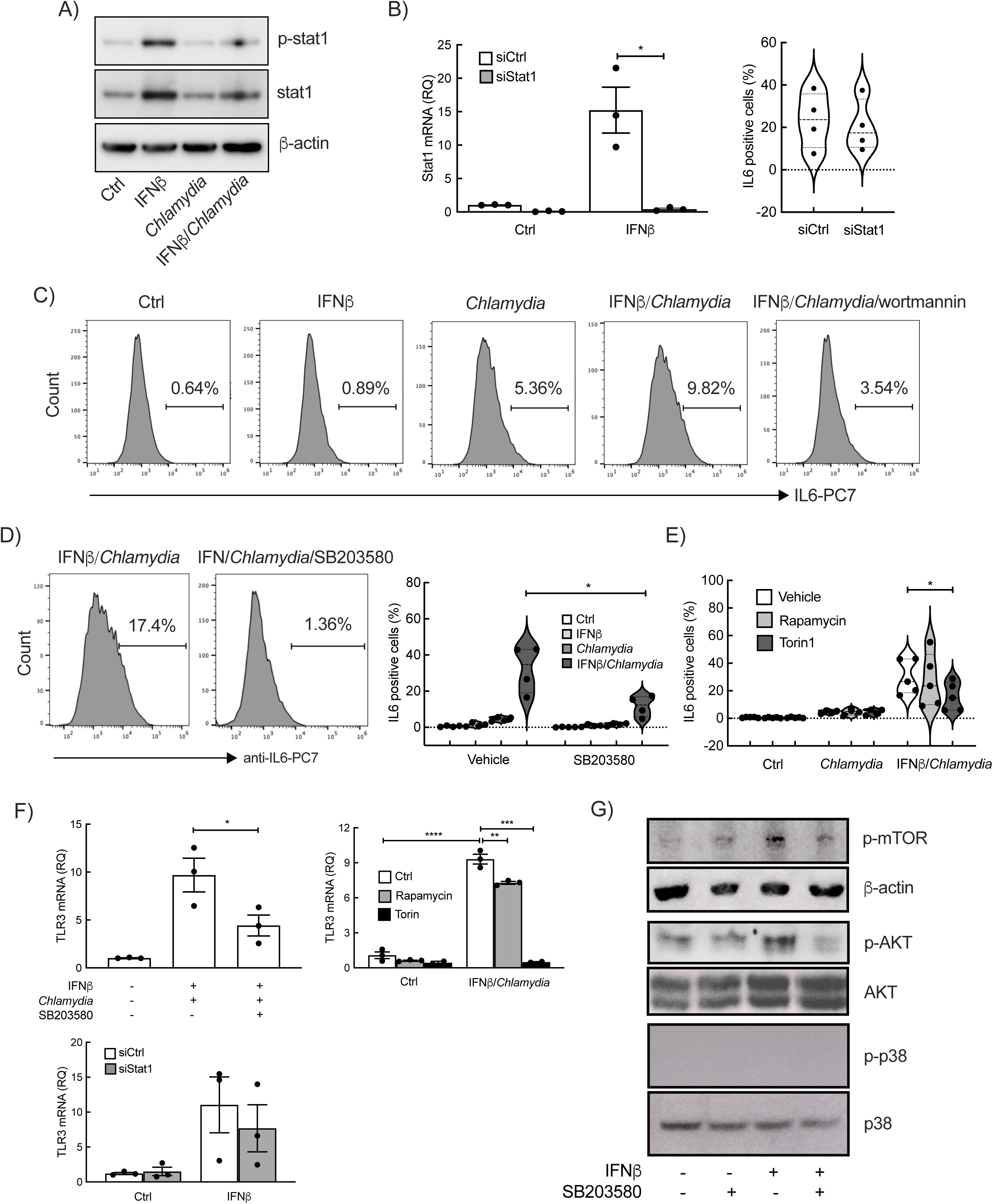
PI3K/AKT-mTOR but not STAT1 signaling pathways downstream of IFN-I are implicated in the synergy between IFN-I and *C. trachomatis*. (A) HeLa cells were incubated with IFNβ and/or *C. trachomatis* for 24 h followed by the detection of activation and expression of STAT1 via immunoblot. The results are representatives of three independent experiments. (B) siRNA against STAT1 or irrelevant oligonucleotides were transfected into HeLa cells for 24 h prior to incubation with IFNβ alone (left graph) or with IFNβ and *C. trachomatis* (right graph). The transcriptional level of STAT1 (left graph) was measured by real-time RT-qPCR and normalized to actin transcript following the 2^-ΔΔCt^ method. The data are presented as relative mRNA levels compared to untreated cells and shown as the mean±SE with individual values of three experiments. P-value of Student’s unpaired t-test is shown (* for p < 0.05). Intracellular IL6 protein (right panel) was determined by flow cytometry using anti-human IL6-PC7 antibody. The results of four independent experiments are shown. C-D) HeLa cells were pre-incubated with wortmannin (5 μM) (C) or SB203580 (10 μM) (D) for 1 h. Cells were then treated with IFNβ and/or *C. trachomatis* for 24 h in the presence of these inhibitors prior to brefeldin A addition for 6 h. Intracellular IL6 expression was analyzed by flow cytometry using anti-human IL6-PC7 antibody. The histograms are the representatives of two (C) or four experiments (D), respectively. The results of the four independent experiments are shown as a violin plot in (D), with p-value of a Student’s unpaired t-test (* for p< 0.05). (E) Same as in D using mTOR inhibitors, rapamycin and torin1 (1 μM). The results of five independent experiments and p-value of a Student’s unpaired t-test are shown (* for p< 0.05). (F) HeLa cells were treated with pharmacological inhibitors (upper panels) or transfected with siRNA (lower panel) as described above, followed by IFNβ and/or *C. trachomatis* treatment for 24 h. TLR3 transcripts was measured by rea-time RT-qPCR as above. Data from three independent experiments and p-values of Student’s unpaired t-test are shown (* p< 0.05, ** p<0.01, *** p<0.001 and **** p<0.0001). (G) The cells were incubated with SB203580 (10 μM) for 1 h before addition of IFNβ for 30 min. Phosphorylation of mTOR, AKT and p38 was detected by immunoblot. The results are representatives of three independent experiments.

IFN can also signal via the uncanonical PI3K/AKT/p38 pathway(10). Wortmannin, an irreversible inhibitor of PI3Ks was used to test whether PI3Ks were involved in the synergy between IFN-I and infection. Pre-treatment of cells with wortmannin decreased IL6 production upon infection in the presence of IFN-I (Fig. 3C), indicating that the PI3Ks are part of the implicated signaling cascade(s). SAR405, a specific inhibitor of PI3K/Vps34(37), had no effect (Fig. S3A), suggesting the involvement of class I and/or II PI3K, but not class III PI3K/Vps34. Protein kinase B (PKB or AKT) and MAP kinase p38 lie downstream of PI3Ks. Incubation of cells with SB203580, a commonly used inhibitor of AKT and p38(38, 39), prevented IL6 synthesis upon treatment with IFNβ and *Chlamydia* (Fig. 3D). p38 was not activated by IFN-I (Fig. 3G), nor by the combination of IFNβ and *Chlamydia* (Fig. S3B). In contrast AKT was activated by IFN-I and its phosphorylation was prevented by SB203580 treatment (Fig. 3G). As positive controls, we verified that recombinant human TNFα induced phosphorylation of p38 and AKT, in a SB203580 sensitive manner (Fig. S3B-C). These results suggest that in these cells AKT, but not p38, is involved downstream of IFNβ signaling.

We also explored the implication of the kinase mammalian target of rapamycin (mTOR) in the signaling cascade(s) linking IFN-I and *C. trachomatis* infection to cytokine secretion. Rapamycin, a specific inhibitor of mTOR complex 1 (mTORC1) (40), had no effect on IL6 synthesis (Fig. 3E). In contrast, torin1, which blocks both mTORC1 and mTORC2, markedly decreased IFNβ/*Chlamydia*-induced inflammation measured on transcript (Fig. S3D) and protein (Fig. 3E) levels. These data indicate that mTORC2 is required for the synergy between IFN-I and *C. trachomatis*. mTOR phosphorylation triggered by IFN-I was attenuated in the presence of SB203508, consistent with its position downstream of PI3K/AKT(10) (Fig. 3G).

Altogether, we conclude that the uncanonical PI3K/AKT/mTORC2, but not canonical IFN signaling STAT1, contribute to the exacerbation of inflammation by IFN-I upon *C. trachomatis* infection in epithelial cells.

### Inhibition of PI3K/AKT and mTOR prevented the upregulation of TLR3 expression

If the synergy between infection and IFN-I is mediate by the upregulation of *TLR3* transcription, inhibition of the signaling pathway downstream of IFN-I is expected to hinder the upregulation of *TLR3* transcription. Indeed, pretreatment of epithelial cells with the AKT inhibitor SB203580 attenuated the upregulation of TLR3 (Fig. 3F). A similar attenuation was detected when using Rapamycin, and the effect of Torin1 was even stronger (Fig. 3F). Silencing STAT1 had no effect (Fig. 3F), confirming that the canonical pathway downstream of IFN-I is not involved. These results support the conclusion that the synergy between infection and IFN-I is mediated by the upregulation of TLR3 downstream of PI3K/AKT/mTORC2 signaling.

### The Erk pathway is stimulated downstream of TLR3

We next investigated the signaling intermediates between TLR3 activation and *IL6* expression. TLR3 signaling generally triggers MAPKs, nuclear factor kappa-light-chain-enhancer of activated B cells (NF-ΚB) and interferon regulatory transcription factor 3 (IRF3) activation to induce the expression of inflammatory cytokines(41, 42). NF-ΚB is a transcription factor playing a central role in inflammation. p65, one predominant subunit of NF-ΚB heterodimers, translocates to the nucleus upon activation(43). We tested whether NF-ΚB was activated by *Chlamydia* infection upon IFN-I treatment using HeLa cells expressing a fusion between p65 and the green fluorescent protein. IL1β was used as positive control for nuclear translocation of GFP-p65 (Fig. S4A, left panels). In contrast, GFP-p65 remained cytosolic in *Chlamydia* infected cells(44), even in the presence of IFN-I (Fig. S4A, middle and right panels). These data indicate that NF-ΚB is not implicated in the proinflammatory response to *Chlamydia* infection, even in the synergy with IFN-I. Furthermore, silencing *IRF3* did not abolish the synergy between infection and IFN-I on IL6 expression (Fig. S4B), indicating that the TLR3-IRF3 cascade was also not involved.

*Chlamydia* infection induced phosphorylation of the MAPK/Erk (Fig. 4A), but not p38 (Fig. S3B). IFN-I alone did not activate Erk, but its presence enhanced *Chlamydia*-induced Erk phosphorylation (Fig. 4A). Application of U0126, an inhibitor of MAP/ERK kinase (MEK), reduced Erk phosphorylation induced by *Chlamydia* infection (Fig. 4B). Pre-incubation HeLa cells with this inhibitor also alleviated the synergetic effect of IFN-I and infection on IL6 production (Fig. 4C), without blocking *TLR3* transcription (Fig. S4C). In addition, silencing TLR3 also blocked Erk phosphorylation upon infection (Fig. 4D). Consistently, *Chlamydia*-driven Erk activation was reduced in the TLR3-KO cells (Fig. 4E). Altogether, these data converge to place the activation of the MAPK/Erk downstream of TLR3 activation upon *Chlamydia* infection.

**Fig. 4.**
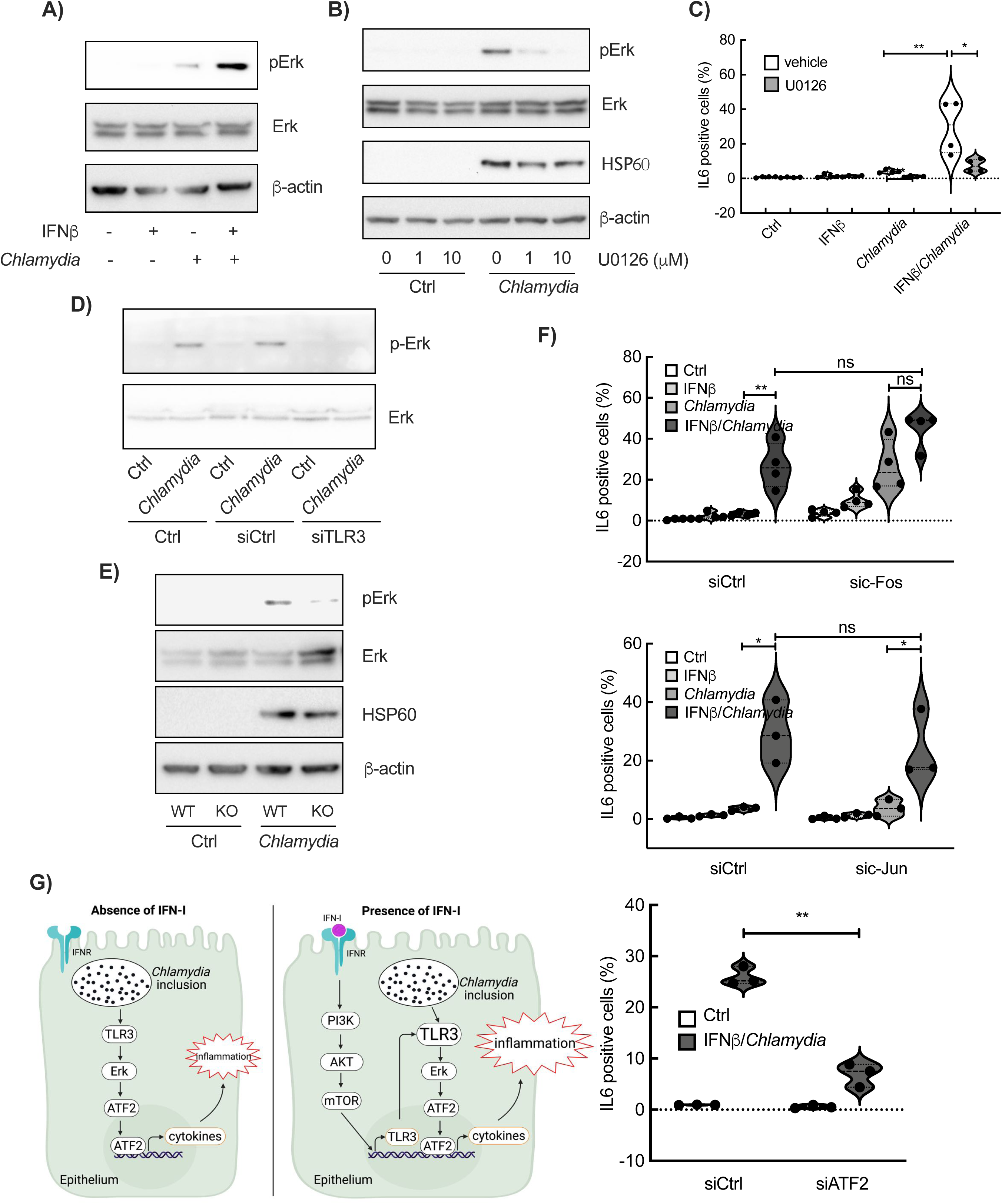
TLR3 modulates the host inflammatory response to the infection by through Erk-ATF2. (A) HeLa cells were treated with IFNβ and/or *C. trachomatis* for 24 h. The activation of Erk and its expression were determined by immunoblot using anti-phosphorylated Erk and anti-Erk antibodies, respectively. The blots are representatives of 3 experiments. (B) The cells were pre-treated with U0126 at the indicated concentrations. One hour later, HeLa cells were infected with *C. trachomatis* for 24 h still in the presence of U0126, followed by the detection of Erk phosphorylation as in (A). (C) After one-hour pre-treatment with U0126 (10 μM), HeLa cells were incubated with IFNβ and/or *C. trachomatis* for 24 h in the presence of U0126, prior to adding brefeldin A for 6 h. Intracellular levels of IL6 were measured by flow cytometry using anti-human IL6-PC7 antibody. The results of four independent experiments and p-value of a Student’s unpaired t-test are shown (* p< 0.05, ** p<0.01). (D) HeLa cells were transfected with control siRNA or siRNA (final concentration 30 nM) against human TLR3 for 24 h before *C. trachomatis* infection for additional 24 h. The activation and expression of Erk was determined as in A. The blots are representative of three experiments. (E) HeLa WT cells and TLR3-KO clone were infected with *C. trachomatis* for 24 h before determining Erk activation and expression. The data are representatives of 3 experiments. (F) siRNA against c-Fos (upper panel), c-Jun (middle panel), ATF2 (bottom panel), or irrelevant oligonucleotides were transfected into HeLa cells for 24 h prior to incubation with IFNβ and/or *C. trachomatis* for 24 h. Intracellular levels of IL6 were measured as in (C). The results of three or four independent experiments and p-value of a Student’s unpaired t-test are shown (* p< 0.05, ** p<0.01) (G) Graphical summary created by BioRender.

Finally, we tested the potential involvement of activator protein 1 (AP-1), downstream of Erk. AP-1 is a transcription factor with a structure of homo-or heterodimer composed of multiple family members including activating transcription factors (ATF), c-Jun and c-Fos (45). We observed that silencing ATF2 significantly reduced IL6 synthesis upon infection in the presence of IFN-I, while silencing c-Jun or c-Fos had no effect (Fig. 4F and S5). These results suggest that the AP-1 transcript factor family member ATF2 controls *IL6* transcription downstream of TLR3-Erk signaling (Fig. 4G).

### dsRNA is detected in Chlamydia infected cells

Double strand RNA (dsRNA) is the only known trigger of TLR3 activation. To determine whether dsRNA is produced during *C. trachomatis* infection we stained infected coverslips with the mouse monoclonal J2 antibody against dsRNA, and with the rabbit polyclonal antibody against the inclusion protein Cap1. dsRNA was detected in the cell cytoplasm and was more abundant in infected cells (Fig. 5 and Fig. S6). To control for the specificity of this staining, cells were incubated before staining with a dsRNA specific RNAse, i.e. RNAse-III. Treatment led to a complete disappearance of the dsRNA-associated signal, while the Cap1 signal remained (Fig. 5A). These data indicate that dsRNA is produced during *Chlamydia* infection and could serve as trigger for TLR3 activation.

**Fig. 5.**
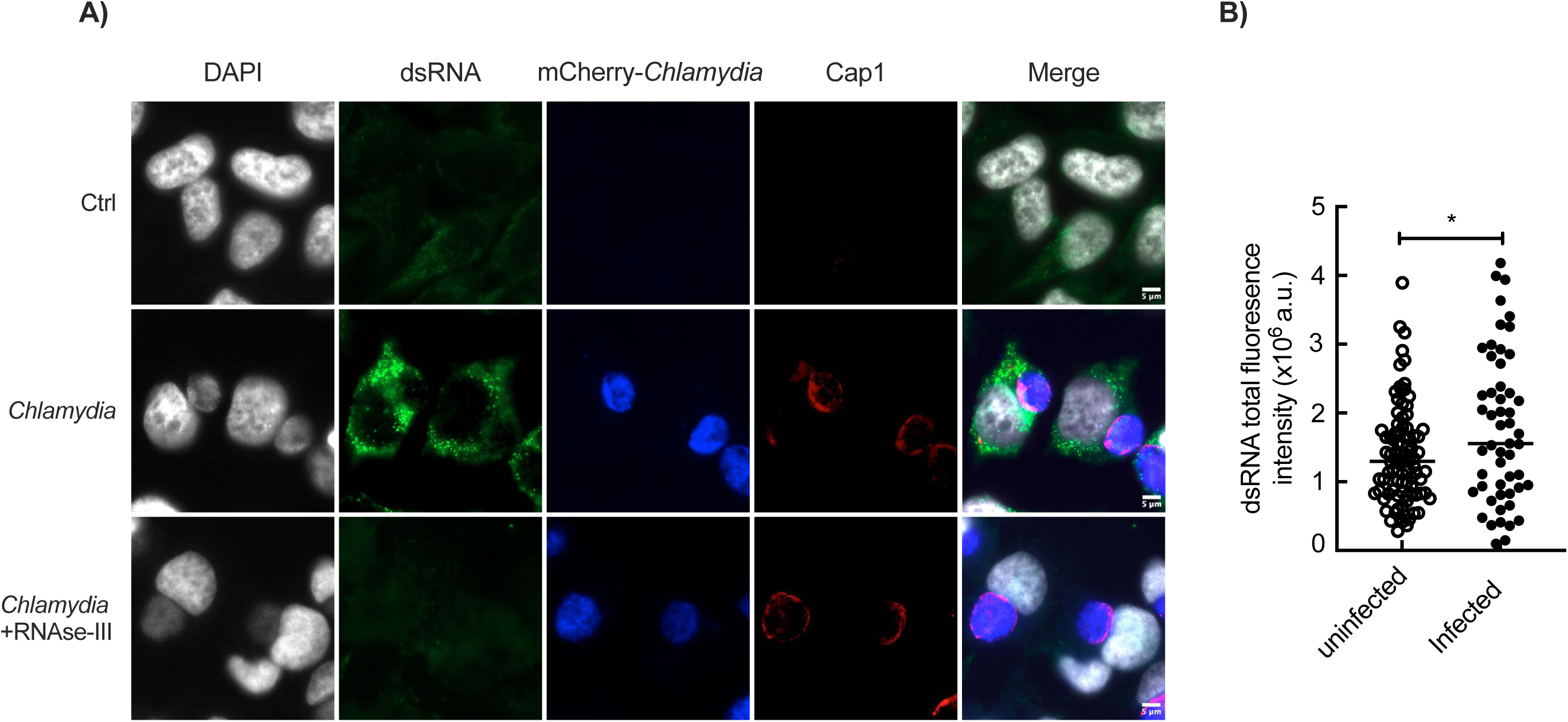
dsRNA accumulates in *Chlamydia*-infected cells. (A) HeLa cells were infected with mCherry-expressing bacteria (MOI=0.3) for 30 h, followed by fixation and immunostaining. DNA was stained with HOECHST 33342 (grey), the inclusion membrane was labeled with an antibody against the bacterial protein Cap1 (red) and the dsRNA was stained with J2 antibody (green). The mCherry signal is displayed in blue. In the lower panels the cells were incubated with RNAse-III for 30 min before immunostaining. The images are representatives of three independent experiments. (B) dsRNA fluorescence intensity in the cytoplasm of infected or non-infected cells was quantified as described in the Methods section and the p-value of a Student’s unpaired t-test is shown (* p< 0.05).

## DISCUSSION

This study examined the effect of IFN-I on the inflammatory response of epithelial cells to *Chlamydia* infection. Treatment of cervical epithelial cells, either primary cells or cancer-derived HeLa cells, with IFN-I strongly increased the synthesis of the pro-inflammatory cytokine IL6 upon *C. trachomatis* infection. This synergetic effect was accounted for by an increase in the expression of the PRR TLR3 upon IFN-I treatment, enhancing *Chlamydia* detection. We determined which signaling cascades, upstream and downstream of TLR3 expression, were implicated.

IFN-I was supplied in the culture medium at nanomolar concentration (2.5 ng/ml or 125 nM), in the range of concentrations detected in the serum of patients with tuberculosis, influenza or COVID-19 infection, ranging from fg/ml to ng/ml(46, 47). Almost all cells in the body synthesize IFN-I(11). Upon *Chlamydia* infection, IFN-I secretion has been detected in cultures of HeLa cells, primary human Sertoli cells of the male reproductive system, and oviduct epithelial cells of mice, indicating that autocrine signaling may occur(19, 21, 23, 48). Consistently, using human fallopian tube derived organoids and human genital serovar of *C. trachomatis* Kessler *et al* reported that *Chlamydia* infection strongly activated the IFN-I cascade pathway (20). IFN-I is not detected in C. *trachomatis*-infected patients, possibly because of its short half-life (approximately 1 to 3 h)(49, 50). Also, the peak of IFN-I induction upon *Chlamydia* infection might be missed, and/or IFN-I synthesis might be local and undetectable in the serum or cervical secretions.

We observed that IFN-I stimulated STAT1 expression, and that this effect was impaired in cells infected with *Chlamydia*, consistent with other studies reporting an inhibition of STAT1 signaling by *Chlamydia* infection(51, 52), and with the absence of effect of STAT1 silencing on the synergy between IFN-I and *Chlamydia* in our study. SB203580 attenuated the synergy between IFN-I and *Chlamydia* as well as the expression of TLR3. Generally, SB203580 inhibits MAPK p38 kinase (IC_50_ ∼ 0.5 μM) and PKB (AKT), another known downstream signal of PI3K (IC_50_ ∼ 5μM)(39). We used SB203580 at 10 μM, which inhibits both kinases. By western blot we detected phosphorylation of AKT but not of p38 upon *Chlamydia* infection, indicating that, in our cellular model, IFN-I promotes TLR3 expression via AKT stimulation.

Using primary human epithelial cells, we observed that IFN-I increased the expression of *TLR2*, *TLR3* and *TLR4*. *TLR3* upregulation by IFN-I was also observed in lung epithelial A549 cells and human umbilical vein endothelial cells (HUVECs) (53). IFNγ also enhanced TLR3 expression in keratinocytes(54). Interestingly, *Chlamydia* infection alone did not trigger the transcription of *TLR3* and showed a tendency to dampen the induction of its transcription upon IFN-I stimulation, indicating that the bacteria may partially counteract IFN-I stimulation (Fig. 2A, S2A). Another study showed that *C. trachomatis* infection elicited TLR3 expression in primary Sertoli cells (48). This increase might rely on IFN-I and an autocrine pathway, as the infection also significantly increased IFNβ production (48). In our model, silencing or knocking-out *TLR3*, but not two other TLRs, attenuated the synergetic inflammation to IFN-I and infection, indicating that TLR3 is a key mediator. Supporting this conclusion, knocking-out *TLR3* has been shown to severely diminish the immune response to *Chlamydia* infection of mouse and human oviduct epithelial cells, including IL6 secretion (3, 55). Interestingly, TLR3 activation itself triggered IFN-I production in murine oviduct epithelial cells infected with *C. muridarum* (56), suggesting a positive feedback loop between TLR3 and IFN.

What activates TLR3 during *C. trachomatis* infection? TLR3 is mainly known for its ability to recognize double-stranded RNA (dsRNA) associated with viral infection.

Recognition of poly(I:C), a synthetic mimic of dsRNA, by TLR3 triggers a strong induction of inflammatory cytokines by macrophages via activation of the NF-ΚB cascade (57). A different signaling cascade is triggered in the context of *Chlamydia* infection as we and others observed a lack of activation of NF-ΚB (44, 58). Here, we show the presence of dsRNA in *Chlamydia*-infected cells. Whether this dsRNA is released by the bacteria or by the host upon *C. trachomatis* infection remains to be determined. Short interfering RNA and mRNA are potential ligand of TLR3, likely through the formation of secondary structures containing stretches of dsRNA (59, 60). Thus, the exact origine and nature of the TLR3 activator during *Chlamydia* infection will need further investigation. To the best of our knowledge, this is the first report of TLR3 activation in epithelial cells by an intracellular bacterium.

Transcription factor AP-1 complexes are key mediators of innate immune responses. The typical heterodimer form of this complex, c-Jun/c-Fos is not involved in TLR3-Erk-mediated inflammatory response to *Chlamydia* infection. In contrast, silencing the other AP-1 component ATF2 significantly decreased the synergic effect between IFN-I and infection. Since silencing c-Jun did not phenocopy ATF-2 silencing, we hypothesize that ATF2 homodimers rather than c-Jun/ATF2 heterodimers(61), might be implicated.

In this study IFN-I was shown to exacerbate *Chlamydia*-induced inflammation in epithelial cells. By exacerbating the pro-inflammatory response of epithelial cells, IFN-I might contribute to the hyper-inflammation experienced by some individuals. Indeed, other studies converge to a deleterious effect of IFN-I production during *C. trachomatis* infection (22, 26–28). The signaling pathways we uncovered can serve as a starting point towards novel therapeutic strategies to alleviate tissue damage after *Chlamydia* infection.

## Acknowledgments

This work was supported by the Institut Pasteur and the CNRS, by the Agence Nationale pour la Recherche 20-PAMR-0011 TheraEPI and the Institut national du cancer INCA_16719. CT and XC were supported by the Pasteur - Paris University (PPU) International PhD Program and funded by a stipend from the National Vaccine and Serum Institute (Beijing, China), and from the Lanzhou Institute of Biological Products (Lanzhou, Gansu Province, China), respectively, both subsidiary companies of China National Biotec Group Company Limited. CT received a stipend from the China Scholarship Council (CSC).

## Author contributions

Conceptualization and writing: YW and AS. Investigation: CT, XC, BN, YVL. Image analysis: FL. Data analysis: CT and YW.

---

**Fig. S1.**
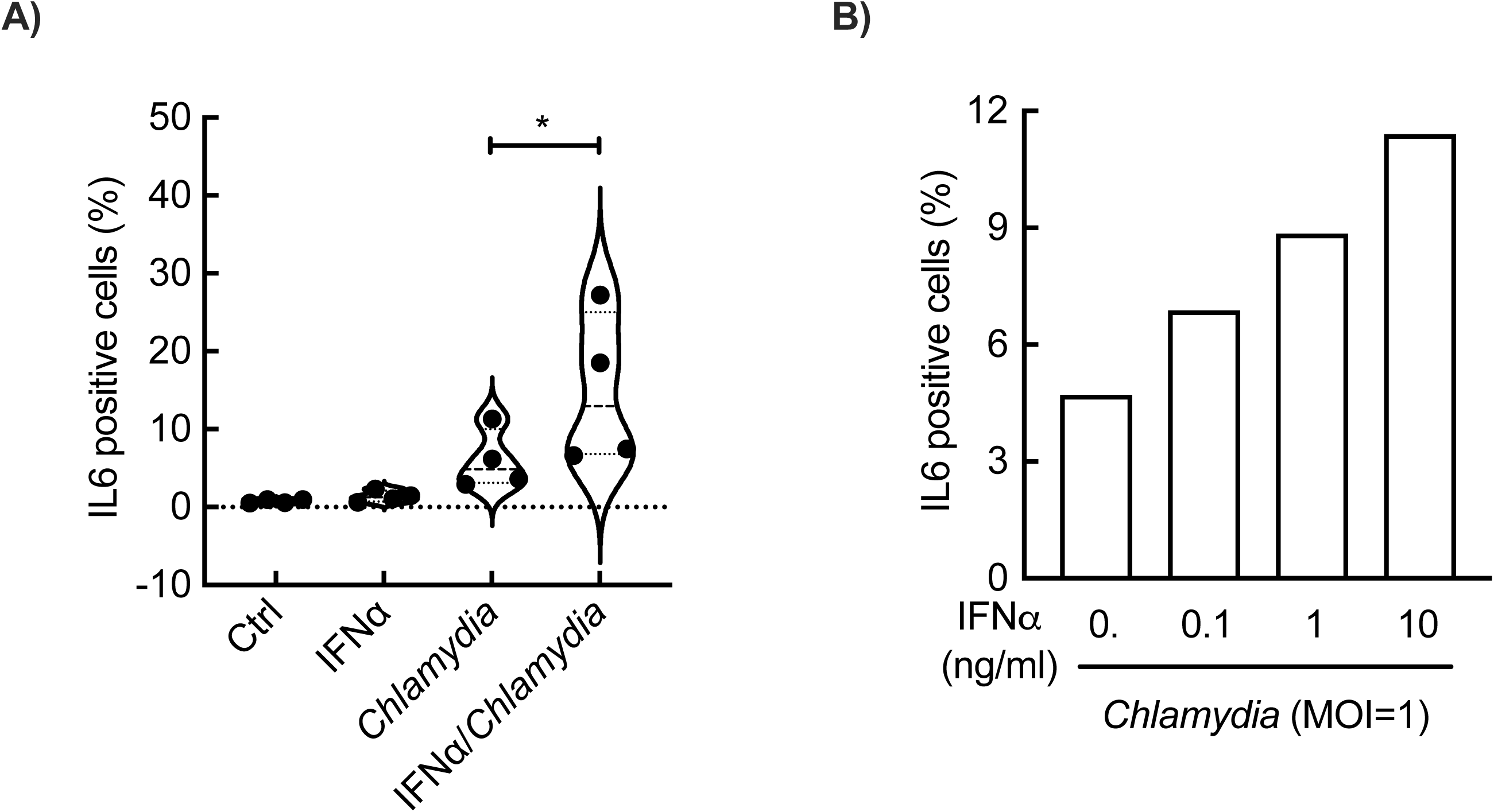
IFNα exacerbates *Chlamydia*-induced inflammation. (A) HeLa cells were incubated with IFNα and/or *C. trachomatis* for 24 h before adding brefeldin A for 6 h. Intracellular IL6 protein was determined by flow cytometry using anti-human IL6-PC7 antibody. The results of four independent experiments and the p-value of a Student’s unpaired t-test are shown (* p<0.05). (B) The same as in A, using IFNα at the indicated concentration. Data are representative of two experiments.

**Fig. S2.**
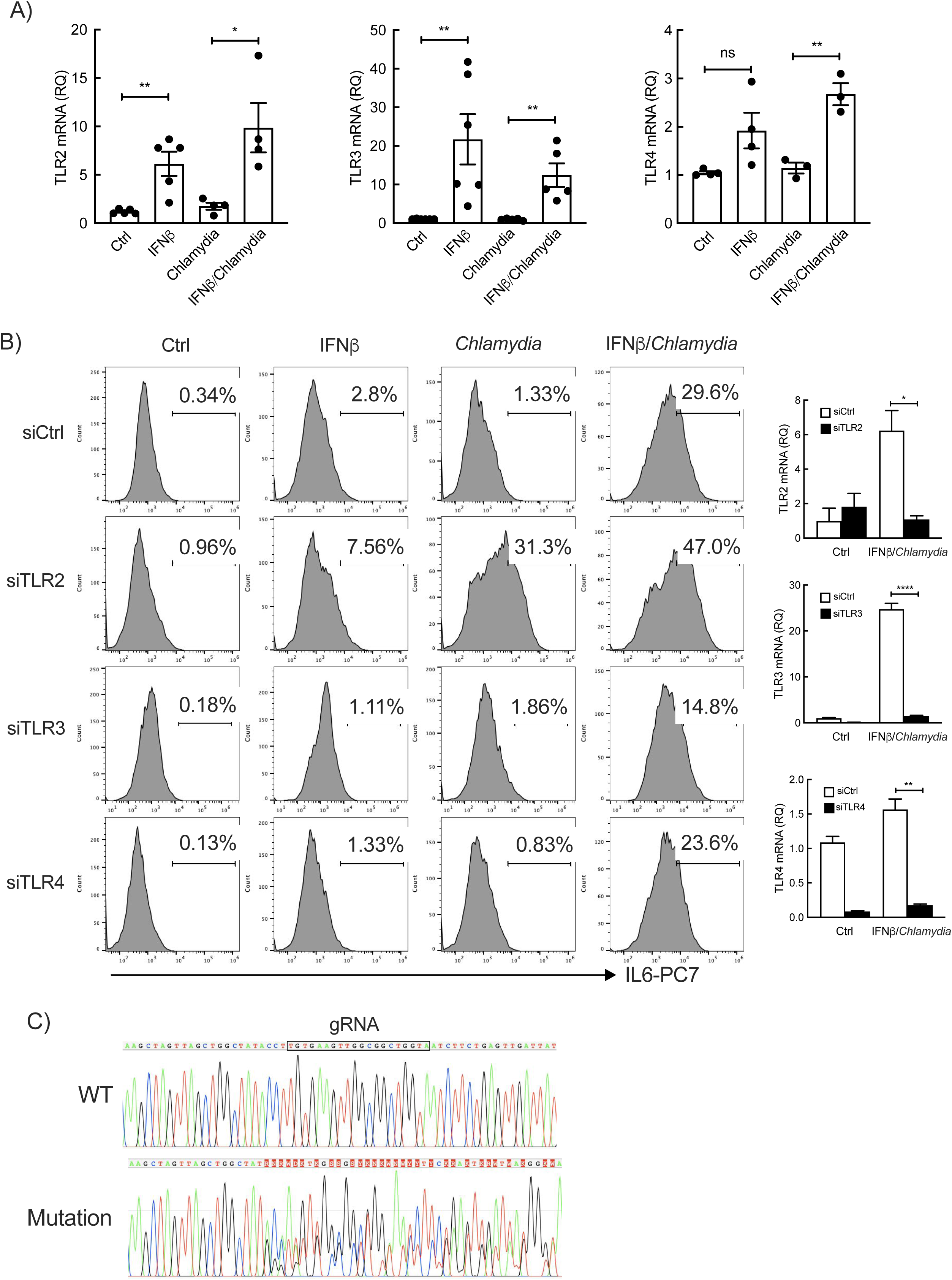
Expression of pattern recognition receptors (PRRs) and their roles in the synergy between IFNβ and *Chlamydia*. (A) HeLa cells were treated with IFNβ and/or *C. trachomatis* for 24 h, followed with RNA extraction. The transcriptional level of *TLR2*, *TLR3* and *TLR4* was determined by rea-time RT-qPCR and normalized to actin transcripts following the 2^-ΔΔCt^ method. The data are presented as relative mRNA levels compared to untreated cells. Each dot represents the result from one experiment. P-values of Student’s unpaired t-test are shown (* p< 0.05 and ** p<0.01). (B) The HeLa cells were incubated with siRNA for 24 h prior to treatment with IFNβ and/or *C. trachomatis* for another 24 h. Brefeldin A was added for 6 h before examining the intracellular IL6 by flow cytometry using anti-human IL6-PC7 antibody. The mRNA levels of the indicated PRRs were determined by quantitative RT-PCR (right panel). The histograms are the representatives of two individual experiments. The RT-PCR experiments are representatives of two independent experiments (right panel). P-values of Student’s unpaired t-test are shown (* for p< 0.05, ** for p<0.01 and **** for p<0.0001). (C) Sequencing data for TLR3-WT and for the knock-out clone generated by CRISPR-Cas9.

**Fig. S3.**
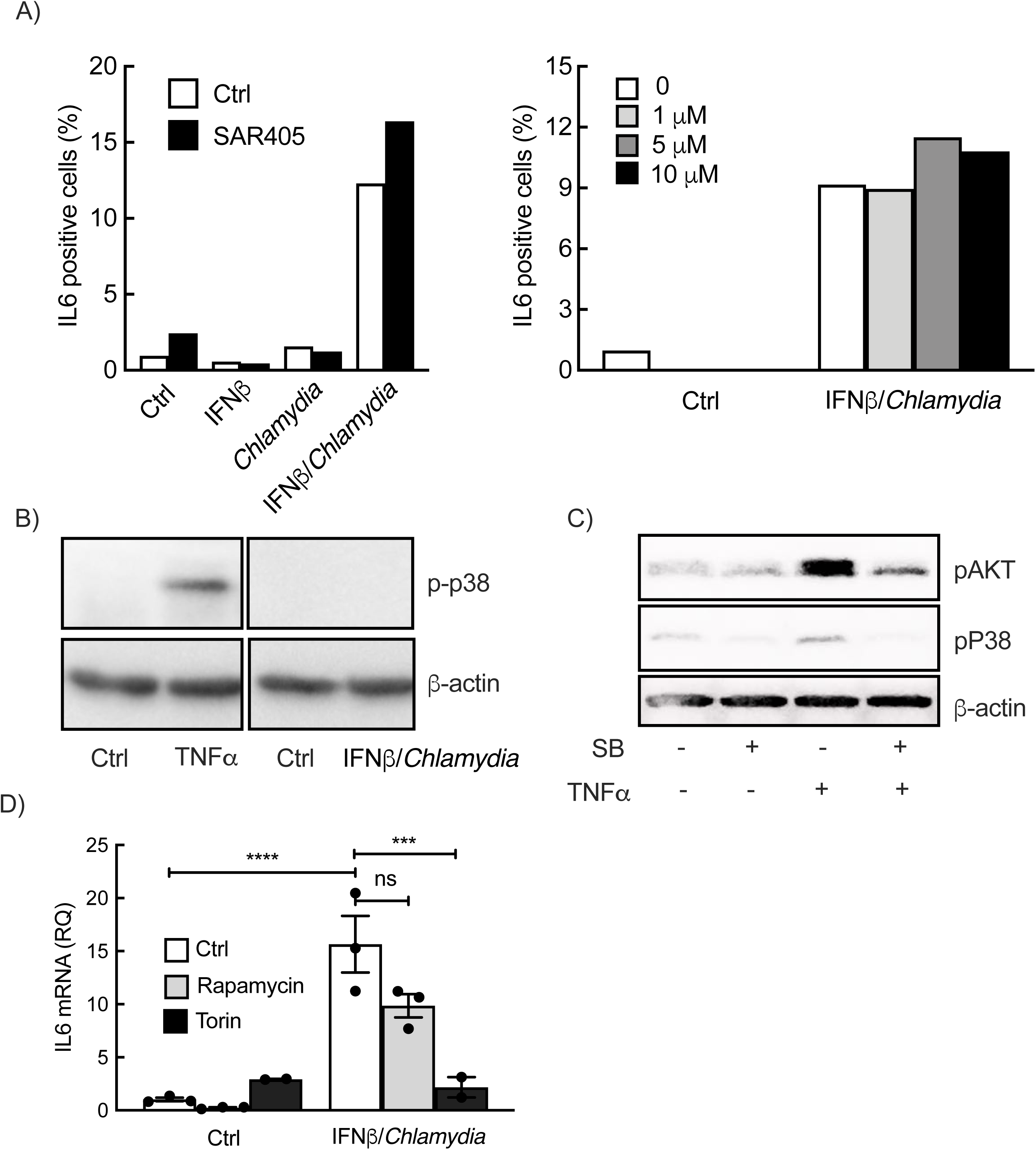
PI3K/Vps34, MAPK/p38 and mTOR complex 1 are not implicated in the synergy between IFN-I and *Chlamydia*. (A) HeLa cells were pre-treated with SAR405 at 400 nM (left panel) or at the indicated concentration (right panel) for 1 h before treating with IFNβ and/or *C. trachomatis* for 24 h in the presence of inhibitor. Brefeldin A was then added for 6 h prior to fixation and measure of IL6 levels by flow cytometry. The data are representative of two individual experiments. (B) HeLa cells were treated with IFNβ and *C. trachomatis*. Twenty-four hours later, phosphorylation of p38 was detected by immunoblotting. As positive control, HeLa cells were incubated with recombinant human TNFα (10 ng/ml) for 30 min followed by p38 phosphorylation detection. (C) HeLa cells were pre-treated with SB203580 (10 μM) for 1 h before incubating with TNFα (10 ng/ml) for 30 min in the presence of SB203580, followed by p38 and AKT phosphorylation detection by immunoblot. The results represent two independent experiments. (D) HeLa cells were pre-incubated with the inhibitors rapamycin (1 μM) or torin1 (1 μM) for 1 h before addition of IFNβ and *C. trachomatis* for 40 h. After treatment, IL6 transcripts were measured by real-time RT-qPCR and normalized to actin transcript following the 2^-ΔΔCt^ method. The data of three independent experiments and p-values of Student’s paired t-test are shown (*** p< 0.001 and **** p<0.0001).

**Fig. S4.**
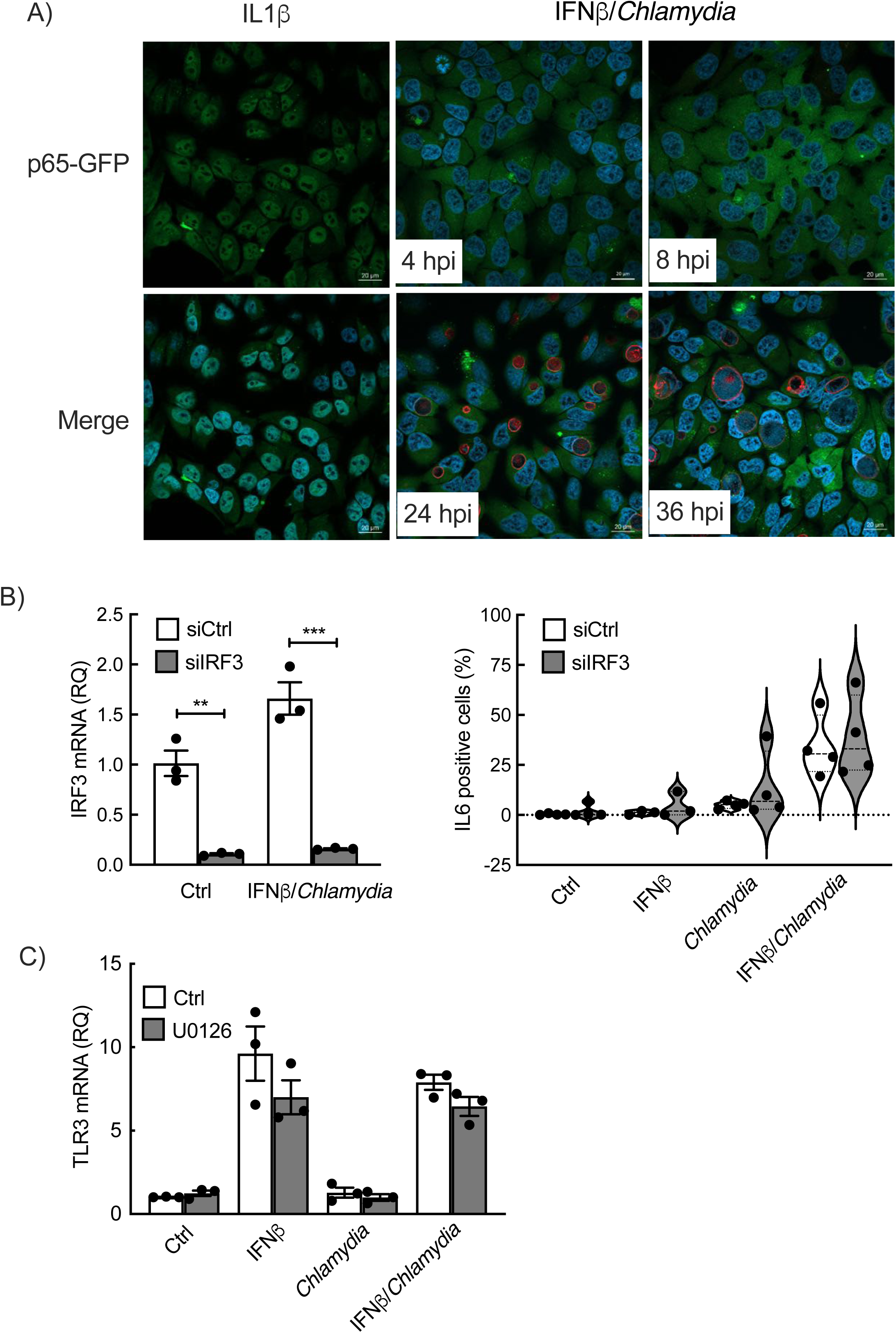
NF-ΚB and IRF3 are not implicated in the synergy between IFNβ and *C. trachomatis* infection. (A) p65-GFP expressing HeLa cells were seeded on coverslip and incubated with recombinant human IL1β (10 ng/ml) for 30 min (left panel), or with IFNβ and *C. trachomatis* for the indicated time (middle & right panels). After cell fixation and permeabilization DNA was stained with Hoechst 33342 (blue) and the inclusion membrane was labeled with an antibody against the bacterial protein Cap1 followed with Alexa647-conjugated secondary antibody (red). The p65-GFP signal is displayed in green. The images are representatives of three experiments. (B) HeLa cells were incubated with siIRF3 or irrelevant oligonucleotide for 24 h. The cells were then treated with IFNβ and *C. trachomatis* for 30 h before analysis of IRF3 transcription by RT-PCR (left panel) and intracellular IL6 by flow cytometry (histograms and violin plot). The result from three (left) and four (right) independent experiments and the p-values of Student’s unpaired t-test are shown (** p<0.01 and *** p<0.001). (C) HeLa cells were pre-incubated with the inhibitor U0126 (10 μM) for 1 h before adding IFNβ and/or *C. trachomatis* for 40 h. TLR3 transcripts were measured by quantitative RT-PCR as described above. The data are presented as mean ± SE of three independent experiments.

**Fig. S5.**
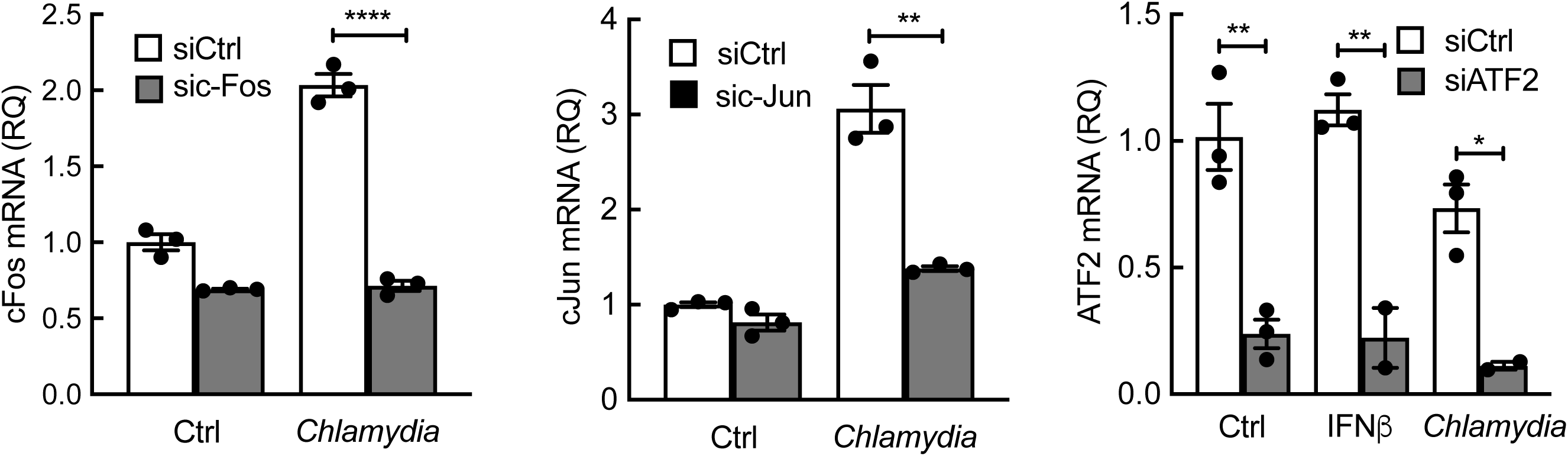
Efficacy of the siRNA at silencing c-Fos, c-Jun and ATF2 in HeLa cells. HeLa cells were transfected with control siRNA or siRNA against human c-Fos, c-Jun or ATF2 (final concentration 30 nM) for 24 h before *C. trachomatis* infection or IFNβ treatment for 24 h. Transcription levels for the indicated genes were measured by real-time RT-qPCR and normalized to actin transcript following the 2^-ΔΔCt^ method. The data of three independent experiments and p-values of Student’s unpaired t-tests are shown (* p< 0.05, ** p<0.01 and **** p<0.0001).

**Fig. S6.**
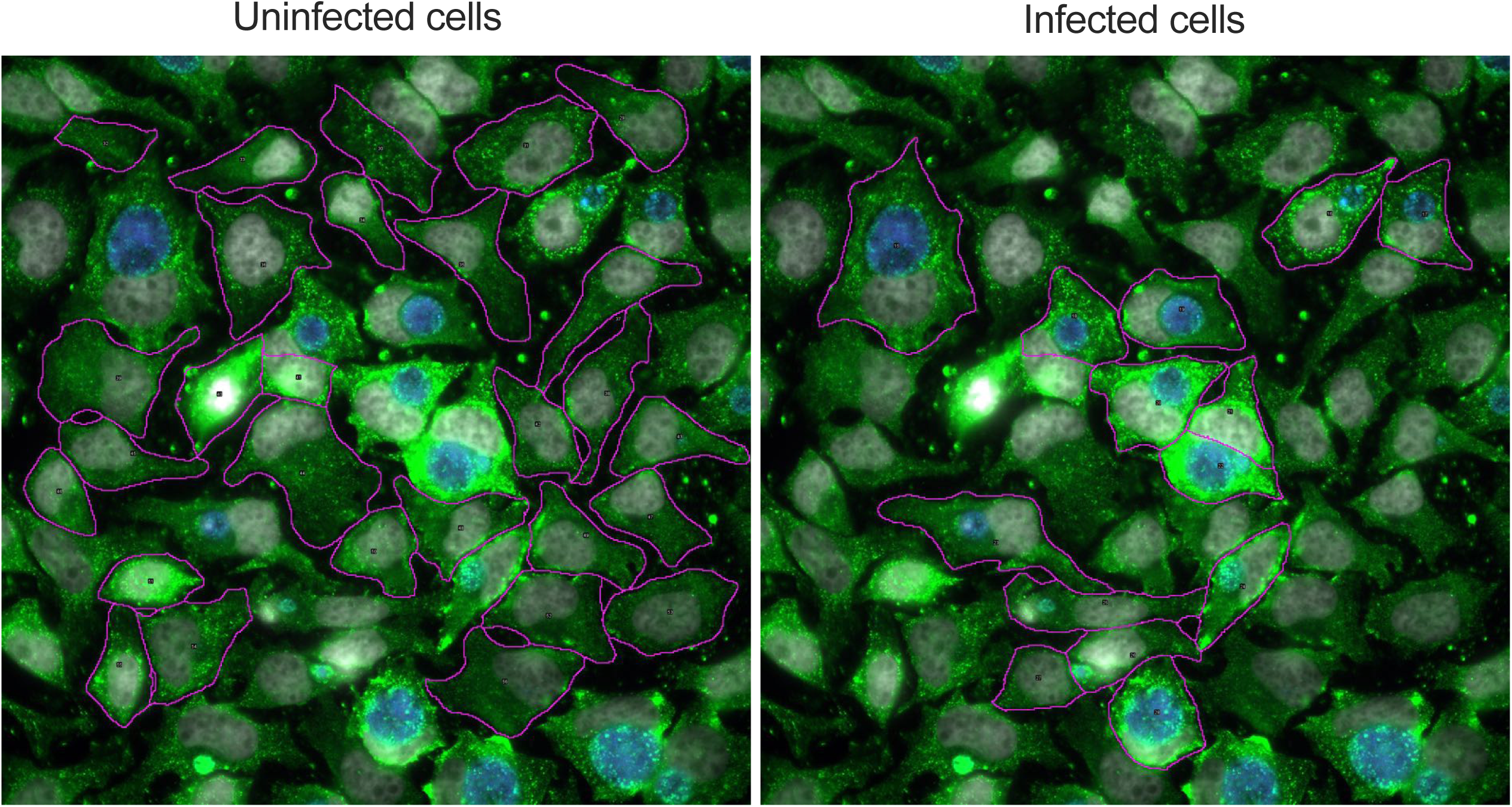
Example of automated detection of “Non-infected” and “Infected” cells. Cells were segmented (purple line) and the presence of an inclusion within the cell identified infected cells. Quantification of the dsRNA signal was performed on the cytoplasm (excluding the signal overlapping with nuclei or inclusions), see Materials and Methods for details.

**Table S1.**
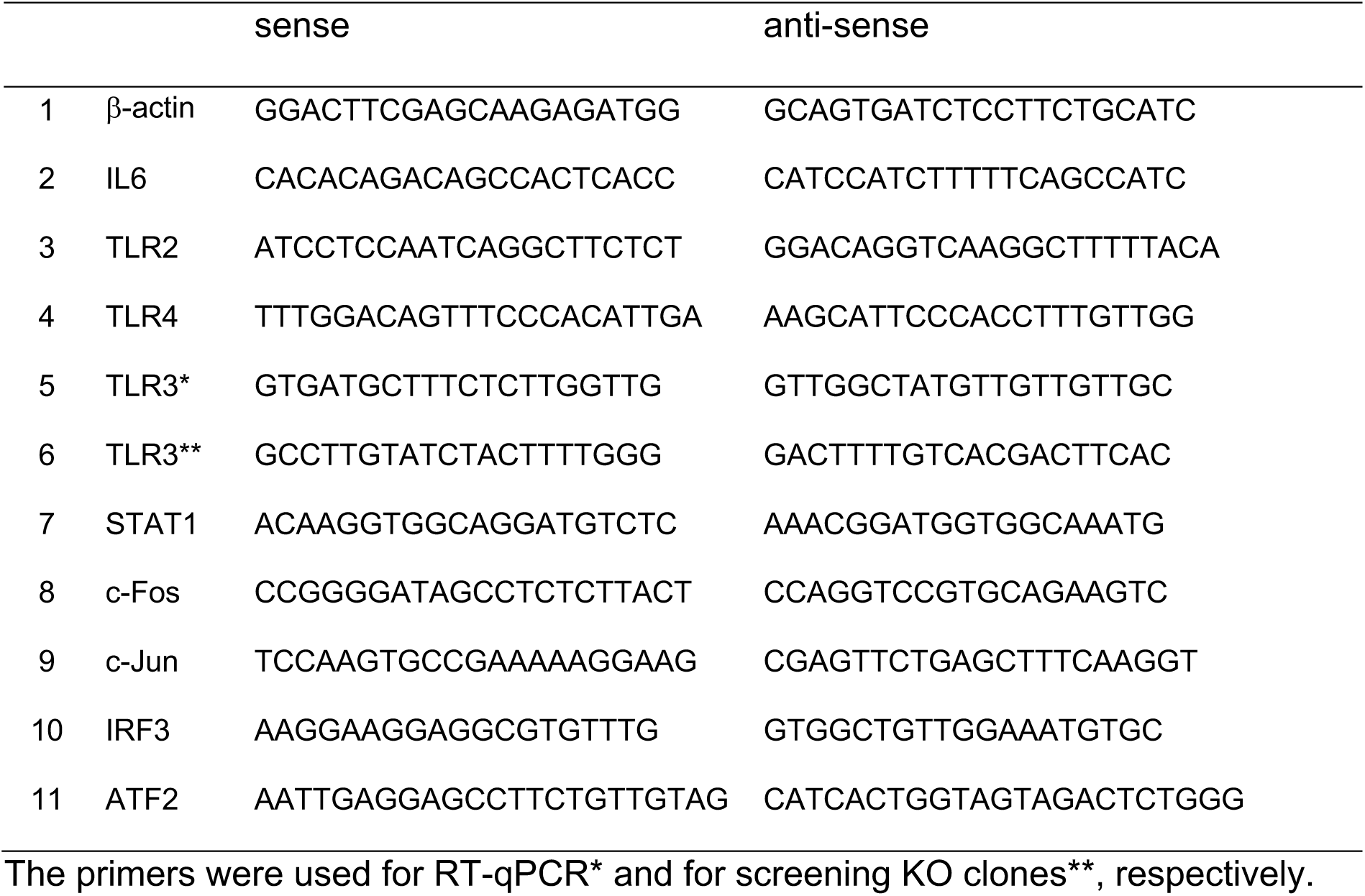
Sequences of the primers used.

